# Endothelial c-IAP2 Loss Amplifies P2X7 Receptor-Driven Inflammation and Worsens Infection-Associated Pulmonary Hypertension

**DOI:** 10.1101/2025.05.22.655387

**Authors:** ES Villarreal, Y Marinho, O Loya, SY Aboagye, DL Williams, J Sun, S Erzurum, V de Jesus Perez, SD Oliveira

**Author notes:** Correspondence: Suellen D. Oliveira, Ph.D.; Assistant Professor - University of Illinois Chicago; College of Medicine; 835 S. Wolcott Ave (M/C 513) Medical Sciences Building Room E713; Chicago, IL, USA - 60612. co-first authorship (ES Villarreal, SD Oliveira).

## Abstract

Schistosomiasis-associated Pulmonary Hypertension (Sch-PH) is the most common form of group I PH worldwide. Recently, data revealed that the preclinical animal model of Sch-PH exhibited gut and lung microbiome dysbiosis linked to significant endothelial dysfunction and microvascular apoptosis, but the role of pro/anti-apoptosis sensors, such as the inhibitor of apoptosis protein 2 (c-IAP2) and purinergic receptor P2X7 (P2X7R), remained unclear. Using a novel Cdh5cre-ER^T2^;cIAP1^-/-^;cIAP2^fl/fl^ animal model, this study investigated the contribution of endothelial c-IAP2 in this process, revealing P2X7R overexpression as a putative target in the onset of Sch-PH. Pharmacologically, inhibition of P2X7R function confirmed its role in promoting lung endothelial death and disease progression. Moreover, data suggest that microbiome-associated metabolic alterations in Sch-PH seem linked to microvascular endothelial apoptosis driven by ATP/P2X7R overactivation and suppressed c-IAP2 expression. Indeed, genetic ablation of endothelial c-IAP2 expression was sufficient to induce PH-like features in mice, with echocardiography indicating a higher pulmonary acceleration time (PAT), PAT/pulmonary ejection time (PET), and right ventricular free wall thickness after IP/IV-Egg challenge compared to controls. These findings suggest a significant contribution of lung endothelial P2X7R activation and c-IAP2 suppression to Sch-PH pathology, highlighting them as promising novel therapeutic targets for this life-threatening illness.

## I. INTRODUCTION

Pulmonary arterial hypertension (PAH) is a life-threatening disease with no cure characterized by injury and hyperproliferation of lung vascular cells, including endothelial cells (ECs).^1^ The hyperproliferation of “abnormal” vascular cells promotes remodeling of pulmonary arteries and arterioles, eventually leading to irreversible inflammatory lesions that collectively drive the pulmonary pressure to life-threatening levels.^2,3^ Although the primary trigger of non-infectious PAH appears multifactorial, several studies indicate it results from a chronic inflammatory process, such as that triggered by infection with the intravascular parasite *Schistosoma mansoni*.^4–6^ After infecting the definitive human or rodent host, *S. mansoni* migrates through the entire cardiovascular system, reaching the mesenteric circulation where the parasite lays its eggs. Within the mesentery, the eggs either cross the intestinal wall, disturbing the gut microbiome, or migrate to other organs, including the lungs, where they can lead to disruption of the lung microbiome homeostasis and cause PAH.^7^ Affecting over 200 million people, it is estimated that *S. mansoni* infection results in schistosomiasis-associated PAH (Sch-PAH) in approximately 1–20 million individuals, making the disease the leading cause of PAH globally.^8,9^ Sch-PAH also shares key pathological features with other forms of PAH, such as the idiopathic disease, including TGF-β-driven severe pulmonary vascular remodeling and obliteration.^10,11^ These changes significantly increase pulmonary vascular resistance, leading to right ventricular (RV) failure and eventual death. Despite its significant global impact and yet unclear epidemiological data in non-endemic countries, schistosomiasis remains in the realm of neglected diseases, with no targeted therapies or specific biomarkers of Sch-PAH early onset.^12^

In general, Sch-PAH progression involves a biphasic inflammatory response. During the acute phase of *S. mansoni* infection, a type 1 helper T-cell (Th1) response promotes pathogen killing but also contributes to tissue damage. As the infection persists and thousands of eggs are released, a type 2 helper T-cell (Th2) response dominates, driving tissue fibrosis, excessive vascular cell proliferation, and severe vascular remodeling characteristic of PAH.^6,10^ This aligns with one long-standing paradigm in the PAH research field that suggests chronic vascular injury leads to a subset of lung ECs to overcome programmed apoptotic cell death, reprogram, proliferate, and thus contribute to the development of inflamed vascular lesions.^13^ Our previous data indicated that during experimental and idiopathic PAH-induced fibrotic vascular remodeling, a subset of lung ECs deficient in expression of the endoprotective proteins Caveolin-1 (Cav-1) and bone morphogenetic protein receptor 2 (BMPR2) survived, proliferated, and contributed to vascular remodeling by stimulating the secretion of pro-fibrotic TGF-β from macrophages.^14^ Similarly, our recent findings indicated that exposure to antigenic *S. mansoni* eggs promoted extensive apoptosis of lung microvasculature and severe inflammatory remodeling, also associated with depletion of lung EC-BMPR2 and Cav-1 expression.^7^ As a scaffold protein and major component of the caveolae, Cav-1 expression and phosphorylation modulate several signaling pathways systemically and within the lung tissue, including death/survival-linked molecular mechanisms, such as the expression of the pro-inflammatory and damage-associated receptor of the purinergic family known as P2X7 receptor (P2X7R).^15^

P2X7R has been previously implicated in experimental schistosomiasis and a non-infectious model of pulmonary hypertension (PH, when referring to animal models).^16–18^ P2X7R activation occurs in response to mM extracellular adenosine-5-triphosphate (eATP) released after cell stress, injury, and/or death.^19^ Persistent P2X7R activation induces the formation of a pannexin pore, leading to a massive influx of Ca^2+^ into the cells, maturation, and secretion of pro-inflammatory cytokines such as IL-1_ and IL-18, and thus, perpetuating inflammation and cell death by inducing pyroptosis and apoptosis.^20^ Moreover, purinergic signaling profoundly affects chronic pulmonary inflammation in response to infectious and “commensal or innate” microorganisms, such as those shaping the gut and lung microbiome.^21–23^ Indeed, gut-lung microbiome dysbiosis, along with cell damage, has been implicated in the release of eATP, supporting the immune response but also favoring microbe colonization.^24^ eATP has also been described as regulating gut microbiota through iron-chelating ability and biofilm dispersion.^21^ In line with these observations, our recently published data demonstrated that the preclinical Sch-PH model displayed gut and lung microbiome dysbiosis, causing significant lung microvascular EC apoptosis by an unclear mechanism.^4,7^

Apoptosis is a complex phenomenon known to be endogenously inhibited by members of the Inhibitors of Apoptosis Protein (IAP) family.^25^ The IAP proteins or <u>B</u>aculovirus IAP repeat-<u>c</u>ontaining proteins (BIRC) gene family regulates apoptotic cell death via cell cycle progression and caspase activation/inhibition.^26^ To date, the IAP family comprises eight proteins known as XIAP, c-IAP1, c-IAP2, NAIP, Livin, Cp-IAP, Op-IAP, and Survivin.^27,28^ All IAP members contain a RING (Really Interesting New Gene) domain, which allows them to function as E3 ubiquitin ligases, catalyzing the ubiquitination process of target molecules and auto-ubiquitination in members such as c-IAP2.^26^ c-IAP1 and c-IAP2 also contain a unique caspase-recruitment domain (CARD), which facilitates interactions such as caspase-8-mediated apoptosis in tumor necrosis factor (TNF)-mediated signaling.^26,29^ c-IAP1 and c-IAP2 also have crucial functions in regulating canonical NF-κB signaling by inhibiting the non-canonical pathway,^30^ but unlike c-IAP1, c-IAP2 expression is inducible. Since there are no specific therapeutic targets for Sch-PH and the mechanisms underlying the disease are poorly understood, our data shed light on the novel molecular contribution of c-IAP2 in the pathogenesis of the disease, given its special auto-ubiquitination and CARD domain attributes. Therefore, we tested the hypothesis that *S. mansoni* egg-induced lung microbiome dysbiosis may contribute to eATP increase within the lung tissue, leading to overactivation of P2X7R-mediated cell death and preventing the rise of inducible anti-apoptotic antagonist c-IAP2, sustaining prolonged lung vascular injury and therefore leading to the inflammatory vascular remodeling that underlies Sch-PH.

## II. MATERIAL AND METHODS

A detailed description of the methods is provided in the Supplementary Information.

### 2.1. Preclinical Animal Model of Sch-PH

The Sch-PH mouse model was performed as previously established by Dr. Brian Graham’s group.^5,10,31^ Briefly, 3-month-old male and female animals were weighed, and then they received an intraperitoneal (IP) sensitization using 240 *S. mansoni* eggs per gram of animal body weight (bw), followed by intravenous (IV) tail injection with 175 eggs per gram/bw two weeks later. ^5,32^ Tail vein injection was performed in 2.5% isoflurane-anesthetized mice, and the depth of anesthesia was monitored by lack of response to toe pinch. For dilation of the tail veins, the mouse tail was submerged in 37°C water or warmed with a heating lamp (held ∼ 20 cm away to avoid overheating or burning) for 20-30 seconds before injection using a 30G needle of the 100 µl of sterile phosphate buffer solution (PBS) alone or containing the eggs. Mice were monitored daily, and after 7 days (Day 21 - IP/IV Eggs established), animals were anesthetized using ketamine/Xylazine (K/X @ 100 and 10 mg/kg body weight; IP). Mice were randomized by sex for subsequent experiments and had *ad libitum* access to food and water. Strain- and age-matched mice were used, as approved by the Institutional Animal Care and Use Committee.

### 2.2. *S. mansoni* cycle and egg collection

All animal studies at Rush University Medical Center were approved by the Institutional Animal Care and Use Committee of the Rush University Medical Center (Department of Health and Human Services animal welfare assurance number A-3120 − 01) with protocol ID: 24-057.

*Biomphalaria glabrata*, strain NMRI, infected with *S*. *mansoni*, strain NMRI, was provided by the NIAID Schistosomiasis Resource Center for distribution through BEI Resources (contract HHSN272201000005I). Three-week-old female Swiss-Webster mice from Charles River were housed in the Comparative Research Center of Rush University Medical Center. As previously described, mice were infected by percutaneous tail exposure to about 200 *S*. *mansoni* through natural transdermal penetration of the cercariae for one hour.^33,34^ Seven weeks post-infection, *S. mansoni* eggs were collected from the isolated liver using standard methods.^33^ Then, eggs were counted, resuspended in sterile PBS, and transported in a sealed container for subsequent exposure to mice.

### 2.3. Novel Conditional Endothelial Cell-Specific BIRC3 Knockout Strain

To investigate the role of both constitutive c-IAP1 and inducible c-IAP2 in the progression of preclinical Sch-PH, we generated a novel animal model of systemic c-IAP1 and conditional endothelial c-IAP2 deletion using *Cdh5* as an endothelial promoter. Specifically, Male and Female 8-12 weeks old heterozygous and homozygous animals (*Cdh5creER^t^*^2^*; c-IAP1^+/-^:c-IAP2^+/fl^*and *Cdh5cre-ER^T^*^2^*;cIAP1^-/-^,cIAP2^fl/fl^*, respectively) were IP injected with corn oil (vehicle) or 1 mg/day tamoxifen for five consecutive days to induce cre-mediated recombination and specific EC-c-IAP2 deletion (after 14 days of the last injection).^35^ After genetic recombination was confirmed by polymerase chain reaction (PCR), Western blot, and immunohistochemistry (IHC) analysis, the Sch-PH model was performed as described above.

### 2.4. Genotyping using Polymerase Chain Reaction

Approximately 1 cm tail samples were collected from adult animals in a sterile environment and used to perform PCR analysis. PCR was performed via a REDExtract-N-AMP^TM^ Tissue PCR kit protocol (Sigma, Cat #XNAT-100Rxn). For the tail digestion, a solution containing 50 µl of extraction solution and 12.5 µl of tissue preparation solution was mixed with individual tail samples and incubated at room temperature for 10 minutes. After the initial incubation, the samples were incubated at 95^°^C for

3 minutes before mixing them with 50 µl of neutralizing solution B. Different genotyping protocols were performed depending on the protocol specificities of the gene of interest. The PCR products were then run in a 2-3% agarose gel electrophoresis, and subsequently, the gels were scanned with Li-Cor Odyssey CLx (Lincoln, NE).

### 2.5. Assessment of right ventricle systolic pressure (RVSP) and Hypertrophy (RVH)

After the animals underwent general anesthesia using K/X as described above, the surgical area was disinfected using 70% alcohol, and a small skin incision was performed to access the jugular vein. Then, a Millar Mikro-Tip catheter transducer (model PVR-1030) was inserted into the RV via the jugular vein to measure RVSP. RVSP was calculated using an MPVS-300 system connected to a Powerlab A/D converter (AD Instruments, Colorado Springs, CO). After recordings, mice were ventilated, and about 1 mL of blood was collected using 3.8% sodium citrate-treated syringes via vena cava puncture. The remaining blood within the lungs was cleared by pump perfusion of 5 mL of cold PBS via a cannula placed in the RV. After complete perfusion, the lung lobes were either snap-frozen in liquid nitrogen or inflated with a 4% paraformaldehyde (PFA) solution for histological analyses. Hearts were dissected for evaluation of RVH using the Fulton index (RV/left ventricle + septum weight ratio), as routinely performed.^7,11^

### 2.6. Experimental Rodent Echocardiography

At day 0 (D0 - baseline) or day 21 (D21) after IV/IP PBS or Egg exposure, heterozygous and homozygous c-IAP2 mice were anesthetized using inhaled isoflurane (2.5% - 3%) and placed in the supine position on a heating pad. Then, individual animals were subjected to transthoracic echocardiography using Vevo F2 (VisualSonics Inc., Toronto, ON, Canada) and a UHF57x transducer. A rectal probe continuously monitored body temperature (T = 36.5-37.5 °C).

### 2.7. Histological analysis of Pulmonary Vascular Remodeling and IHC

PFA-fixed, paraffin-embedded lung sections (5 μm) were used to evaluate protein expression. Fluorescent images were collected using an LSM880 confocal microscope (Carl Zeiss MicroImaging, Inc.). In addition, histological analysis of microvessel area and thickness was quantified in *Masson’s Trichrome-*stained sections.^3^

### 2.8. *In situ* Apoptosis Assay

*In situ* apoptosis was performed as previously published,^7^ using terminal deoxynucleotide transferase (TdT)-mediated dUTP nick-end labeling (TUNEL) detection kit (ab206386; Abcam; Massachusetts, US) according to the manufacturer’s instructions. The ratio of TUNEL-positive to total cells (apoptotic index) was measured within the lung microvasculature (vessels > 100 μm).

### 2.9. Competitive Enzyme-Linked Immunosorbent Assay

Competitive Enzyme-Linked Immunosorbent Assay (ELISA) was carried out following the protocol from the mouse baculoviral IAP repeat-containing protein 3 (BIRC3; c-IAP2) ELISA Kit (MyBioSource, Cat #MBS7252940). The optical density was then measured at 450 nm in a microplate reader (Accuris Smartreader 96), and the results were plotted on GraphPad Prism.

### 3. Human Microvascular ECs from lungs (HMVEC-L) and pulmonary artery ECs (HPAEC)

Primary HMVEC-L and HPAEC (4th-7th passage) were obtained from Lonza (Cat No. CC-2527 and CC-2530, respectively), maintained in endothelial basal medium-2 (EBM-2) supplemented with endothelial growth medium (EGM-2) SingleQuots^TM^ Supplements (CC-4176; HPAEC) or MV SingleQuots (CC-4147; HMVEC-L) plus heat-inactivated fetal bovine serum up to 10% at 37 °C and 5% CO_2_, following Lonza’s standard operational protocol. Then, 90-100% confluence cells were treated for 18 h with 10-30 ng/mL TNF-_ (Cat No. GF023; Millipore) or 50-100 ng/mL INF-_ (Cat No. IF002; Millipore) and cell lysates used to test the role of inflammatory cytokines on P2X7R expression by western blot. Cell morphology was assessed daily by brightfield contrast microscopy.

#### 3.1. Apoptosis assay by flow cytometry

HMVEC-L seeded in 6-well plates were treated with 3mM ATP, 30 ng/mL TNF-_ (Cat No. GF023; Millipore) with or not 50 µM A740003 (Cat. No. 3701, Tocris) or 200 nM Staurosporine (STS; Cat No. 1285; Tocris) for 18 hrs. The cells were washed with PBS and gently detached using trypsin-EDTA 1× (Cat No. 15400-054; Gibco). Then, 5 × 10^6^ cells were incubated with APC Annexin V Apoptosis Detection kit with PI, according to the manufacturer’s protocol (Cat No. 640932; Biolegend). Data were acquired using Gallius (Beckman Coulter, USA), and the events were quantified using Kaluza 2.2 (Beckman Coulter, USA).

#### 3.2. Sample Preparation and Western Blot

Frozen lung tissue and cultured ECs were fully homogenized using cold radioimmunoprecipitation assay (RIPA) buffer containing 1% protease and 0.1% phosphatase inhibitor cocktail. After western blot, membranes were scanned with Li-Cor Odyssey CLx (Lincoln, NE), and data were analyzed and normalized to β-actin or GAPDH loading controls using ImageJ software (https://imagej.nih.gov/ij/).

#### 3.3. Liquid chromatography-mass spectrometry (LC-MS)

Anti-inflammatory short-chain fatty acids (SCFAs) metabolites were isolated from plasma. Briefly, 5 μL of the calibrator and sample were injected into an AB SCIEX 5500 QTRAP coupled with an Agilent 1290 UPLC system (samples eluted by Agilent Poroshell column 120 EC-C18 2.7 μm, 2.1 x 100 mm with a flow rate of 450 μl/min; column compartment at 40 °C). LC elution was 99% mobile phase A (0.1% FA in H_2_O) for 1 min, followed by a linear gradient increase of mobile phase B (0.1% FA in ACN) from 1% to 10% in 1 min, then from 10% B to 65% B in 6 min and from 65% B to 90% B in 0.1 min. The column was washed with 90% B (3 min), then re-equilibrated to the initial condition (99% A; 3 min). The autosampler was maintained at 4 °C. MS data were acquired by MRM scan in negative mode. The ESI spray voltage and source temperature were held at -4.5 kV and 450 °C. Negatively charged analytes and standards were detected by monitoring their transition to signature product ions.

#### 3.4. Statistics analysis

Data were analyzed using One Codex Cloud Platform and GraphPad Prism v10 (GraphPad, La Jolla, CA, USA). Normally distributed data are presented as the arithmetic mean +/- Standard Error of the Mean (SEM). Shapiro-Wilk test was used to determine the normality of data, and the Brown-Forsythe or F test was used to assess the equality of variances. Then, a parametric or nonparametric test was performed accordingly. Parametric statistical analysis was performed using unpaired Student t-test between 2 groups. One-way or Two-way ANOVA, followed by post hoc analysis (Bonferroni, Dunnett, or Tukey Multiple Comparison tests), was used to analyze differences between > 2 groups. Nonparametric analysis was performed using the Mann-Whitney test. All results were subjected to the ROUT outliers test and were two-sided. P < 0.05 was considered statistically significant.

## III. RESULTS

### 3.1. Preclinical Sch-PH model displays increased expression of lung microvascular endothelial P2X7R

Our recent findings in the preclinical animal model of Sch-PH revealed gut and lung microbiome dysbiosis, along with significant lung microvascular apoptosis, though the underlying mechanism remained unclear.^4,7^ Dysfunctional gut and lung microbiome composition can significantly dampen the circulating level of anti-inflammatory microbial metabolites such as the SCFAs, whereas it can also increase levels of damage-associated molecular patterns (DAMPs), such as ATP, contributing to a sustained inflammatory response. Therefore, to evaluate the contribution of circulating SCFAs on the development of Sch-PH, we first compared plasma acetate, propionate, isovalerate, and butyrate from preclinical IP/IV Egg-exposed animals with vehicle control animals **(Fig. 1A, B)**. MassSpec analysis of plasma SCFAs from IP/IV Egg-exposed and vehicle control animals indicated no significant differences in the level of acetate, propionate, isovalerate, and butyrate between groups **(Suppl Fig. 1A)**. Further functional pathway analysis of gut-derived metadata^7^ using MetaCYC also indicated no significant functional differences in the gut microbiome-linked processes of Sch-PH group compared to controls **(Suppl Fig. 1B)**. On the other hand, similar analysis of lung-derived metadata, pointed to reduced aerobic respiration (cytochrome C) in the IP/IV Egg-exposed group compared to controls **(Suppl Fig. 1C)**, known to potentially change eATP level via cellular stress and injury. Microbiome dysbiosis has already been implicated in contributing to eATP/P2X7R-mediated inflammatory signaling.^23^ In line with these observations, western blot analysis revealed a significant increase in P2X7R expression in the lung tissue of the Sch-PH model, specifically in the microvasculature, compared to larger arterial vessels **(Fig. 1C, D)**. Further *in vitro* experiments using pro-inflammatory stimulation with TNF-U and INF-γ confirmed a selective inflammation-induced microvascular P2X7R expression in the pulmonary circulation. Specifically, TNF-U, but not INF-γ treatment, significantly increased P2X7R expression only in human lung microvasculature (HMVEC-L) compared to human pulmonary artery ECs (HPAEC) **(Fig. 1E-F)**. Overall, the data indicated that *S. mansoni* egg-induced inflammation preferentially increases P2X7R expression in the pulmonary microvasculature during Sch-PH, where high levels of apoptosis were observed.^7^

**Figure 1:**
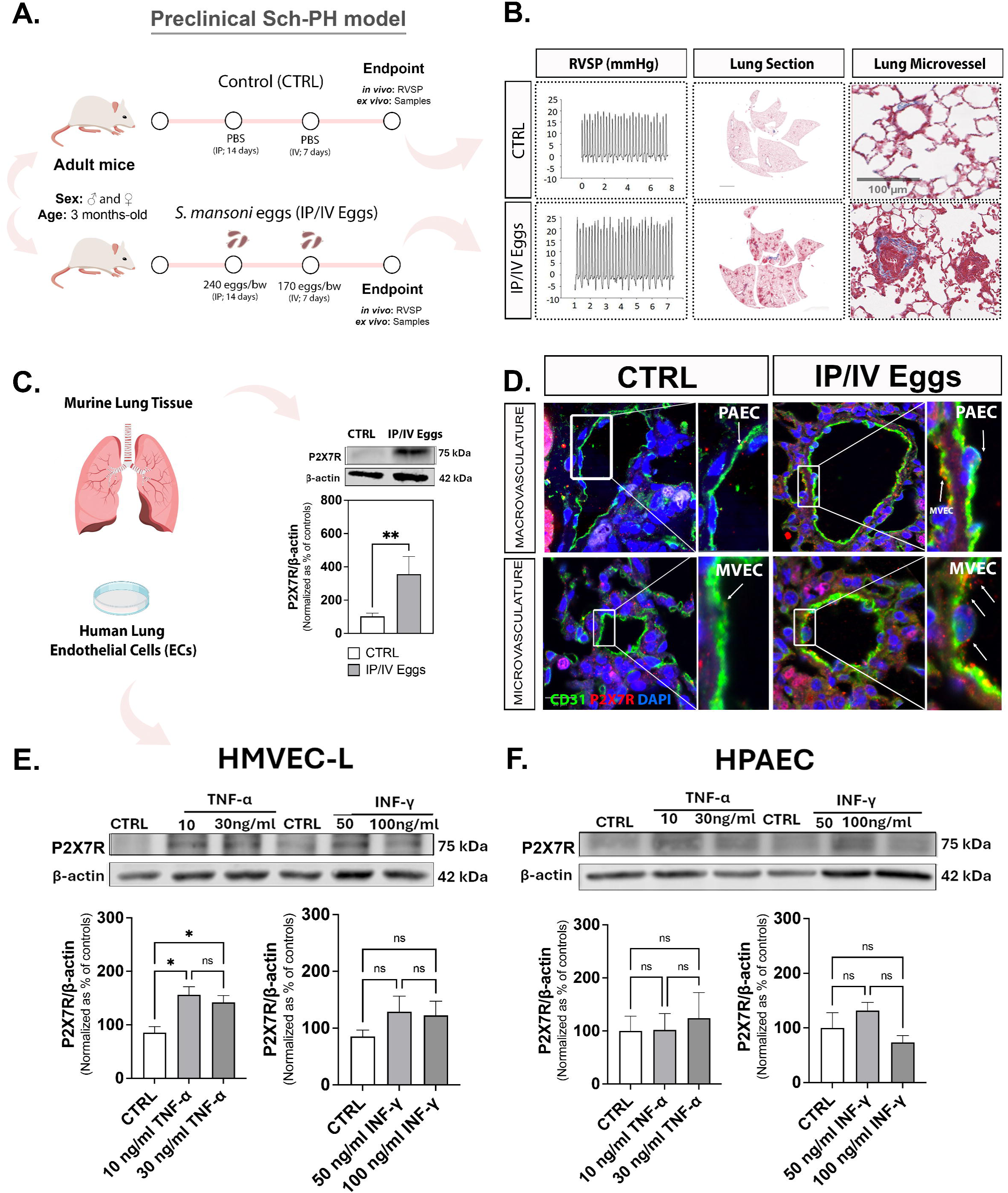
Preclinical Sch-PH model displays increased pulmonary P2X7R expression, including in microvascular ECs. Sch-PH mouse model was induced in 3-month-old mice by intraperitoneal (IP) sensitization using 240 *S. mansoni* eggs/gram of body weight (bw), followed by intravenous (IV) tail injection with 175 eggs/gram bw after two weeks (IP/IV Eggs). Then, after 7 days, male and female animals were used for in vivo hemodynamic analysis (RVSP = right ventricular systolic pressure in mmHg) and tissue procurement for histological analysis of vascular remodeling in the whole lung tissue section using *Masson’s Trichome* staining **(A, B)**. Representative western blot and quantification of the P2X7 receptor (P2X7R) expression in the whole murine lung lysates **(C)**. Representative lung sections from vehicle controls (CTRL) and IP/IV Egg-exposed mice showing P2X7R (red) and CD31 (green) expression in the lung macro and microvasculature; DAPI was used to stain nuclei (blue) **(D)**. Representative Western blot and quantification of P2X7R expression (normalized as % of controls) in control (CTRL), TNF-U (10 and 30 ng/ml), and INF-_ (50 and 100 ng/ml) stimulated lung human microvascular endothelial cells (HMVEC-L; **E**) and human pulmonary artery EC (HPAEC; **F**) (18h). Normally distributed data was analyzed using Student *t*-test or one-way ANOVA (n = 8-10 animals/group; n = three different cultures; ns = non-significant; *P < 0.05; ** P < 0.01).

### 3.2. Increased lung P2X7R and reduced c-IAP2 contribute to lung microvascular EC apoptosis in Sch-PH

ATP-induced P2X7R activation leads to massive influx of Ca^2+^, cytokine secretion, and cell death.^20^ *In vitro*, pharmacological inhibition of P2X7R function in HMVEC-L was sufficient to prevent TNF-U/eATP-mediated late apoptosis **(Fig. 2A-B)**. P2X7R is known to co-localize with Cav-1 in healthy lung vasculature.^36–38^ Reduced lung EC-Cav-1 has been demonstrated to contribute to lung EC apoptosis in PAH,^7^ including by playing a role in the regulation of the anti-apoptotic IAP member, survivin;^39^ however, its impact on the expression pattern of other IAP members, especially in the context of pathogen-linked PH, remains largely unclear. An initial analysis of IAP genes in isolated lung ECs from *Cav1*^-/-^ mice found upregulation of c-IAP2 (BIRC2) and survivin (BIRC5) expression compared to WT cells **(Suppl Fig. 2A)**. In contrast to c-IAP2, survivin lacks the RING and CARD domains, which regulate protein auto-ubiquitination, turnover, and their ubiquitin ligase activity making inducible c-IAP2 unique.^27,28^ However, despite increased mRNA levels, lung tissue of global Cav-1 knockout (KO) and EC-Cav-1 knockout (ECKO) mice were deficient in c-IAP2 expression **(Fig. 2D),** similar to what is observed in the lungs from Sch-PH mice compared to vehicle controls **(Fig. 2E)**. Structurally similar to c-IAP2, the expression of constitutive c-IAP1 remained unchanged between groups **(Fig. 2F)**. Moreover, measurement of HRP-conjugated c-IAP2 level using a competitive ELISA, indicated its reduction in plasma samples from Sch-PH mice compared to controls **(Fig. 2G).** As an inversely proportional assay, it revealed a higher c-IAP2 level in the plasma of Sch-PH mice. Altogether, these data indicate lung c-IAP2 expression is potentially protective against the development of Sch-PH, and its depletion in the lung tissue is in part due to release into the plasma as a response to egg exposure.

**Figure 2:**
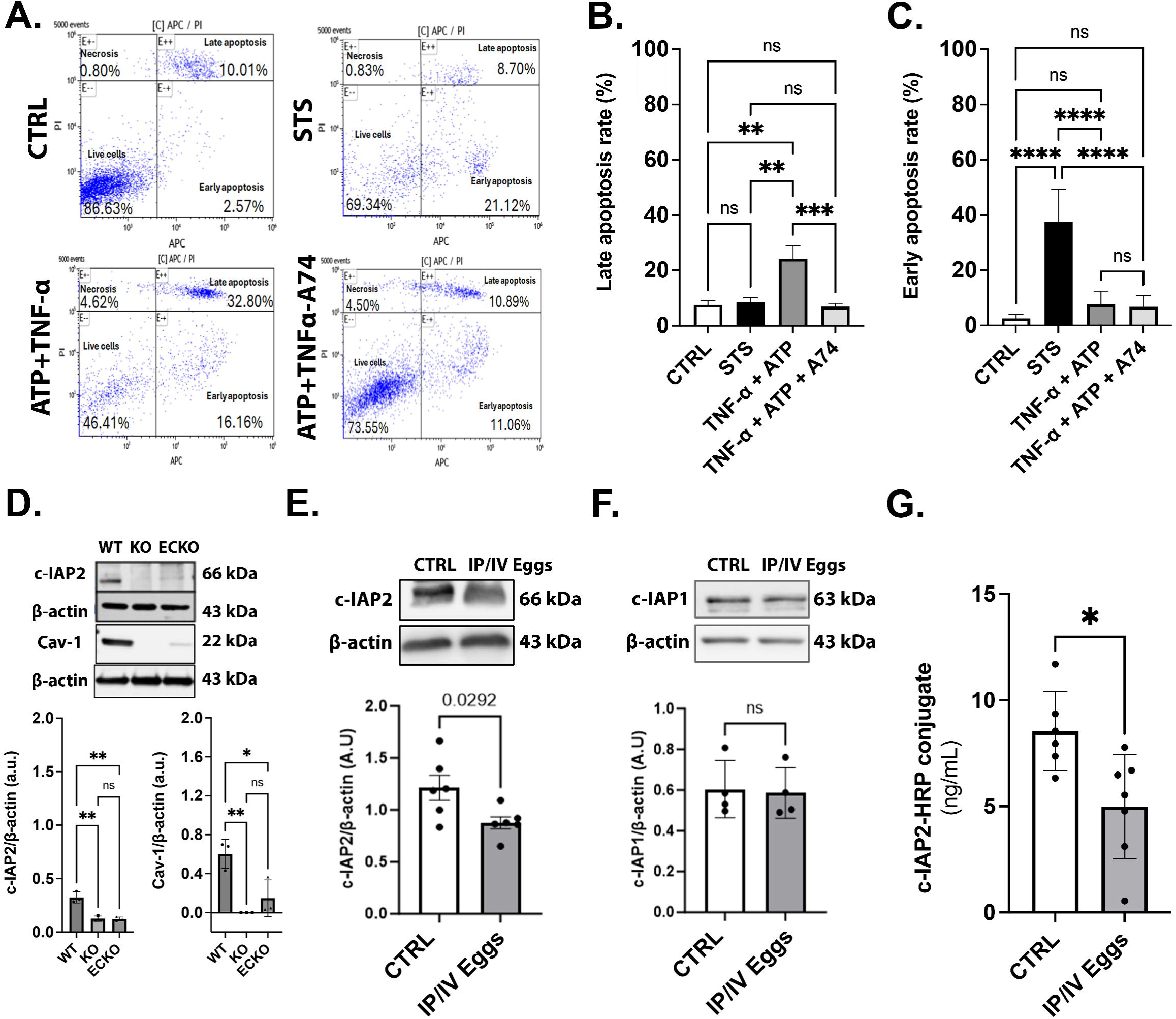
Increased P2X7R and reduced inducible c-IAP2 expression may contribute to lung EC apoptosis in Sch-PH. P2X7R-mediated apoptosis in HMVEC-L after 18 hours of treatment with vehicle control (CTRL) or 30 ng/ml TNF-U plus 3 mM ATP with or without P2X7R antagonist 50 μM A740003 (A74). Staurosporine (STS, 200 nM) was used as a positive control (**A-C;** n= six different cultures). Western Blot analysis of c-IAP2 and Caveolin-1 (Cav-1) expression in whole lung lysates of wildtype (WT), global Cav-1 knockouts (KO), and endothelial-specific Cav-1 knockouts (ECKO) **(D).** Western Blot analysis of c-IAP1 and c-IAP2 expression in whole lung lysates from CTRL and IP/IV Egg-exposed mice **(E, F).** c-IAP2 competitive ELISA using plasma from CTRL and IP/IV Egg-exposed mice (inversely proportional). Normally distributed data was analyzed using Student t-test or one-way ANOVA (n = 3-6 animals/group; ns = non-significant; *P < 0.05; ** P < 0.01; *** P < 0.001; **** P < 0.0001).

### 3.3. Genetic ablation of EC-c-IAP2 increases lung P2X7R expression and leads to spontaneous vascular remodeling and PH in mice

To further investigate the contribution of c-IAP2 expression to EC lung vascular homeostasis, we generated a novel endothelial-specific heterozygous (HET; *Cdh5creER^t^*^2^*;c-IAP1^+/-^:c-IAP2^+/fl^*) and homozygous (HOMO; *Cdh5creER^t^*^2^*; c-IAP1^-/-^:c-IAP2^fl/fl^*) mouse strain under transcriptional control by *Cdh5creER^t^*^2^. Genetic ablation of *Birc2* and *Birc3* and consequent suppression of c-IAP1 and c-IAP2 expression was validated through PCR, IHC, and Western blot analysis **(Fig. 3A, B; Suppl Fig. 2B)**. In addition, no significant difference in lung Cav-1 and phosphorylated-Cav-1 expression was observed among WT, HOMO, or HET animals **(Suppl Fig. 2D, E)**, suggesting lung endothelial-c-IAP2 genetic ablation does not significantly contribute to EC-Cav-1 expression level or phosphorylation in murine lungs. In terms of vascular structural homeostasis and PH, genetic ablation of EC-c-IAP2 expression induced spontaneous remodeling in the pulmonary circulation and a significant increase in RVSP and RVH **(Fig. 3C-G)**, indicating that endothelial c-IAP2 depletion is sufficient to induce experimental PH-like features. Additionally, endothelial c-IAP2 deletion increased P2X7R expression in the whole lung tissue compared with controls **(Fig. 3E)**. Together, these data strongly suggest that genetic ablation of EC-c-IAP2 expression contributes to the development of a mild but spontaneous PH phenotype and increased P2X7R expression in the lungs, indicating a putative connection between lung P2X7R/c-IAP2 signaling pathways.

**Figure 3:**
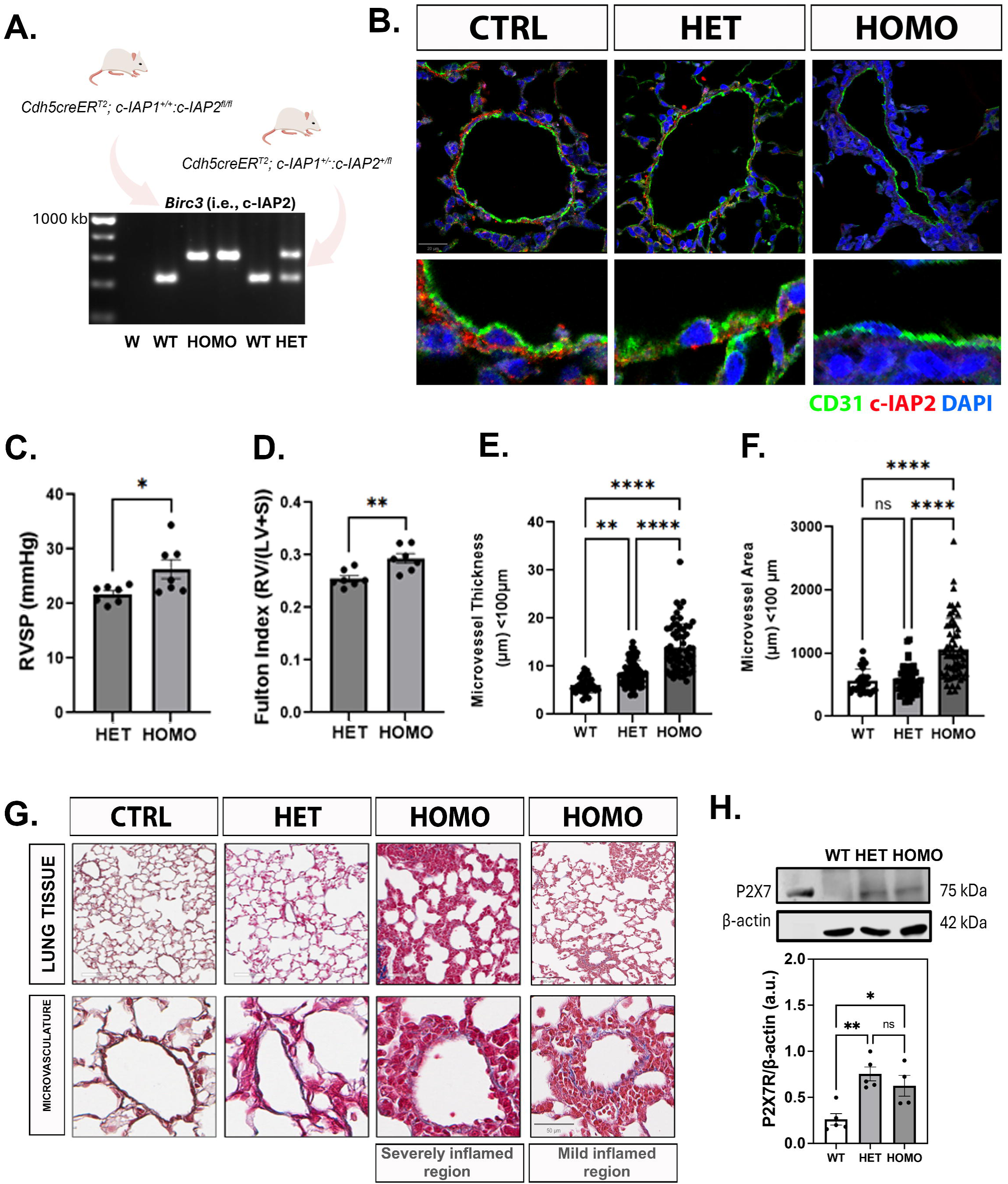
Endothelial-c-IAP2 genetic ablation increases P2X7R expression in the lungs and leads to spontaneous PH. The polymerase chain reaction was used to validate the genetic profile of c-IAP2 (BIRC3) in control (CTRL), heterozygous (HET), and homozygous (HOMO) mouse strain (*Cdh5creER^t^*^2^*; c-IAP1^+/-^:c-IAP2^+/fl^*and *Cdh5cre-ER^T^*^2^*;cIAP1^-/-^,cIAP2^fl/fl^*, respectively) **(A)**. Immunohistochemistry analysis of CD31 (green) and c-IAP2 (red) expression in lung sections from CTRL, HET, and HOMO animals DAPI (blue) stained nuclei **(B)**. Right Ventricular Systolic Pressure (RVSP) and Right Ventricular Hypertrophy (RVH) were measured in HET and HOMO animals **(C)**. Microvessel thickness (μm**; E)** and area (μm^2^; **F)** were quantified in lung sections from CTRL, HET, and HOMO stained using *Masson’s Trichome* **(G)**. Scale bar = 100 μm (top panel) and 50 μm (bottom panel) (vessels with a diameter smaller than 100 μm using the ImageScope software). Western Blot analysis of P2X7 receptor expression **(H)**. Normally distributed data was analyzed using Student *t*-test or one-way ANOVA (n = 3-6 animals/group; ns = non-significant; *P < 0.05; ** P < 0.01). = non-significant; *P < 0.05; ** P < 0.01; *** P < 0.001; **** P < 0.0001).

### 3.4. Genetic ablation of EC-c-IAP2 expression exacerbates *S. mansoni* egg-induced PH

To address the relevance of endothelial c-IAP2 expression for the development of Sch-PH, *Cdh5creER^t^*^2^*;c-IAP1^-/-^:c-IAP2^fl/fl^*underwent echocardiography analyses before (baseline) and 21 days after PBS or IP/IV egg exposure **(Fig. 4A)**. Additionally, lung sections were evaluated to quantify the degree of pulmonary vascular remodeling **(Fig. 4B)**. In the context of lung hemodynamics, *S. mansoni* eggs stimulus reduced Pulmonary Acceleration Time (PAT) and the PAT/Pulmonary Ejection Time (PET) ratio compared with controls **(Fig. 4C-E)**, indicating increased pulmonary vascular resistance. No differences in PET alone were observed **(Fig. 4E)**. Echocardiograph analysis also showed that *S. mansoni* egg stimulus increased RV Free Wall Thickness (RVFWT) compared with control animals **(Fig. 4F)**, confirming RVH. No significant differences were observed in fractional shortening **(Fig. 4G)**. Regarding RV function, no significant changes were observed in Tricuspid Annular Plane Systolic Excursion (TAPSE; **Fig. 4H**). No significant changes were observed in cardiac output (CO), ejection fraction (EF), stroke volume (SV), and heart rate (HR) **(Fig. 4I-L).** Together, the data suggest an essential role of EC-c-IAP2 in Sch-PH development.

**Figure 4:**
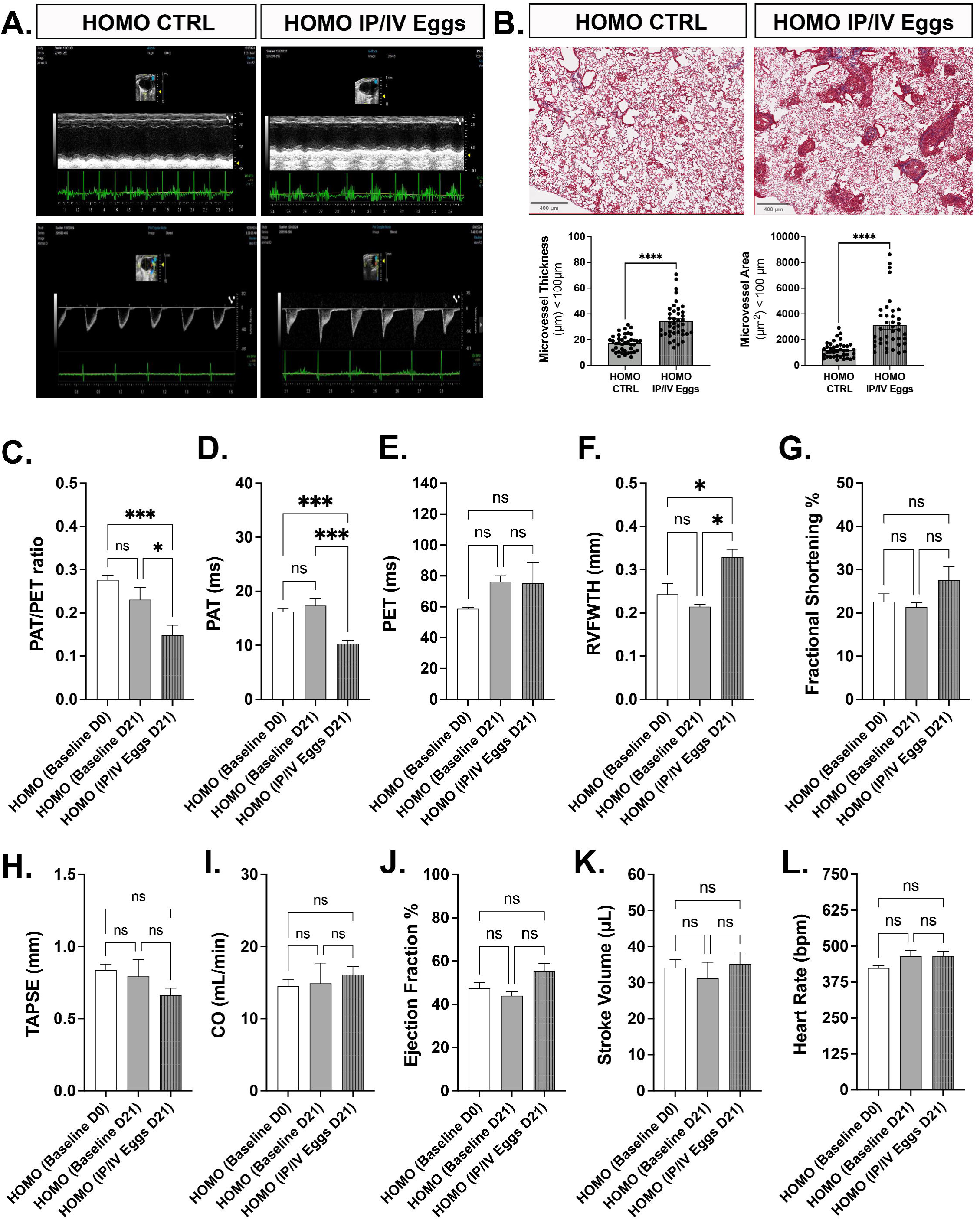
***Schistosoma mansoni* egg exposure induces severe PH in endothelial-specific c-IAP2 knockout mice.** Homozygous *Cdh5cre-ER^T^*^2^*;cIAP1^-/-^,cIAP2^fl/fl^*, (HOMO) mouse underwent echocardiography analysis at day 0 (baseline D0) and day 21 after PBS (baseline D21) or *S. mansoni* egg exposure (IP/IV Eggs D21) (**A**). Microvessel thickness (μm) and area (μm^2^) were quantified in lung sections from HOMO control or IP/IV eggs stained using *Masson’s Trichome* **(B)**. Scale bar = 400 μm (top panel - representative micrograph) and (bottom panel - quantification). Pulse-wave Doppler echo was used to measure pulmonary acceleration time (PAT) and the pulmonary ejection time (PET), and PAT/PET ratio **(A)** or individual PAT **(B)** and PET **(C)** were plotted. **D.** Right Ventricle free wall thickness (RVFWT) was calculated during end-diastole in the parasternal short-axis mitral valve level 2D or parasternal long-axis RV outflow tract level M-mode (inset images at the top). Tricuspid annular plane systolic excursion (TAPSE) was measured in 2D M-mode echocardiograms from the apical 4- chamber view, positioning the cursor on the lateral tricuspid annulus near the free RV wall and aligning it as close as possible to the apex of the heart. Stroke volume (SV), fractional shortening (FS), ejection fraction (EF), and cardiac output (CO) were measured from the left ventricle (LV). Normally distributed data was analyzed using Student *t*-test or one-way ANOVA (n = 3-7 animals/group; ns = non-significant; *P < 0.05; ** P < 0.01; *** P < 0.001).

### 3.5. Pharmacological inhibition of P2X7R ameliorates pulmonary vascular remodeling and prevents RVSP increase in Sch-PH animals but does prevent RVH or fully restore c-IAP2 expression

To test whether P2X7R would serve as a potential pharmacological target to prevent or attenuate Sch-PH in a preclinical setting, animals were IP injected with two doses of 45.5 mg/kg BBG 24 hours before the IP and IV *S. mansoni* egg injections **(Fig. 5A)**. Pharmacological treatment demonstrated efficacy in preventing RVSP increase but showed no significant effect in preventing RVH in the Sch-PH animal model **(Fig. 5B, C)**. BBG treatment also dramatically reduced pulmonary vascular remodeling and the presence of apoptotic cells within the lung tissue of the *S. mansoni* egg-exposed group compared with the animals stimulated only with eggs **(Fig. 5D-G)** but was not able to fully rescue lung Cav-1 or c-IAP2 expression caused by *S. mansoni* egg-exposed mice **(Fig. 5H, I)**. Overall, data revealed P2X7R as a potential pharmacological target to prevent Sch-PH development.

**Figure 5:**
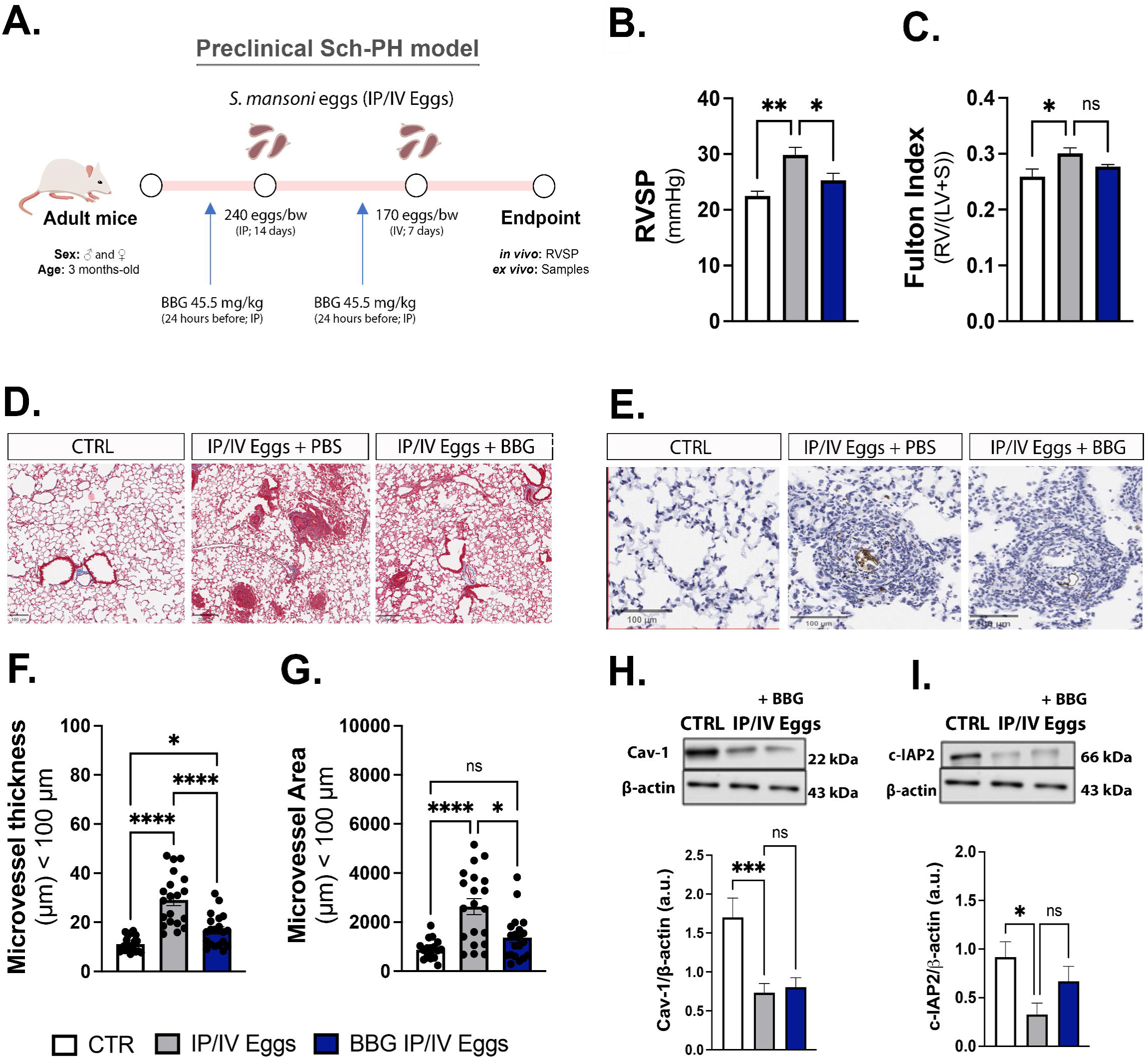
Blockage of P2X7R prevents pulmonary microvascular apoptosis, attenuates right ventricular systolic pressure (RVSP), and remodeling in Sch-PH. Diagrammatic representation of therapeutic approach using two intraperitoneal doses of P2X7R inhibitor brilliant blue G (BBG; 45.5 mg/kg) **(A)**. RVSP **(B)** and RV hypertrophy (RVH; **C**) were quantified in PBS control (CTRL), IP/IV eggs + PBS, and IP/IV eggs + BBG. *Masson’s Trichome* staining (**D**) was used to quantify microvessel thickness (μm**; F)** and area (μm^2^; **G),** and TUNEL analysis to quantify the number of apoptotic cells in lung sections **(E)**. Western blot analysis was performed to quantify lung Caveolin-1 **(J)** and c-IAP2 **(K)** expression. Normally distributed data was analyzed using Student t-test or one-way ANOVA (n = 6-8 animals/group; ns = non-significant; *P < 0.05; ** P < 0.01; *** P < 0.001; **** P < 0.0001).

### 3.5. Sex-linked variations in lung microbiome composition and in the degree of RVH may influence the efficacy of therapeutic inhibition of lung P2X7R in Sch-PH

Sex-linked differences in lung microbiome composition and RVH may significantly impact preclinical drug efficacy by modulating immune response and metabolic pathways. Indeed, our data indicate that only female animals displayed significant RVH in the preclinical Sch-PH model compared to males, which has not been prevented by 45.5 mg/kg BBG treatment **(Fig. 6A, B)**. Moreover, differences in microbial communities between males and females are known to influence the production of key metabolites, such as purine derivatives. Accordingly, lung microbiome composition followed different patterns between sexes, with the overall percentage of the Phylum Firmicutes higher in females compared to males and reduced only in females after egg exposure, an effect not rescued by BBG treatment **(Fig. 6C-F)**. Specifically, the Firmicutes Families Lachnospiraceae, Oscillospiraceae, and Eubacteriaceae displayed reduced readcounts, opposing Lactobacillaceae **(Fig. 6G)**. Thus, taking into consideration changes in the host microbiome composition is essential for optimizing sex-targeted therapies and improving outcomes in pulmonary vascular diseases.

**Figure 6:**
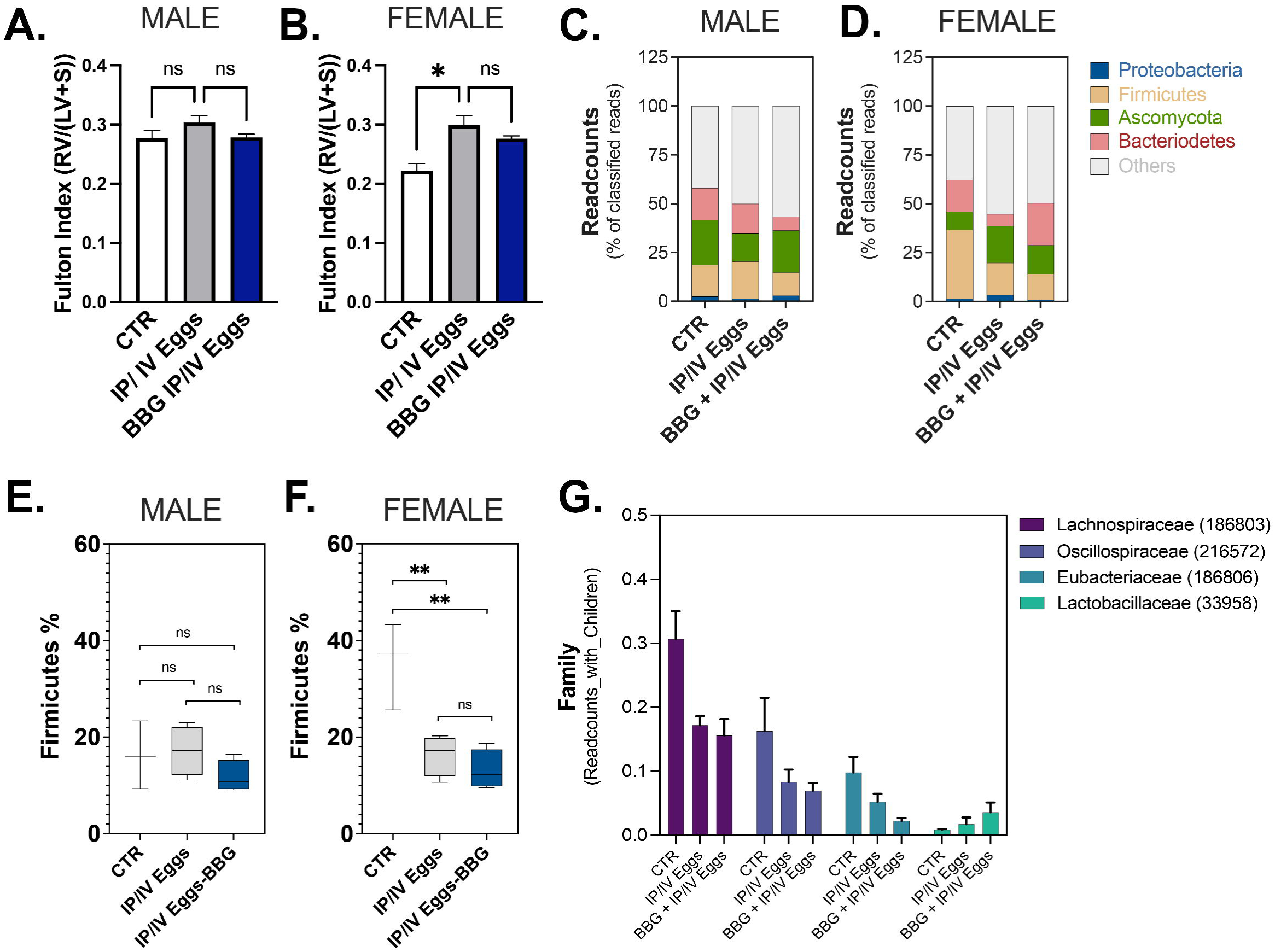
Blockage of P2X7R does not prevent lung microbiome dysbiosis in female Sch-PH. Right ventricle hypertrophy (RVH; **A, B**) and metagenomic analysis **(C-G)** were performed in male and female PBS control (CTRL), IP/IV eggs + PBS, and IP/IV eggs + BBG. Lung tissue was used for whole-genome sequencing and metagenomic analysis using the One Codex Database (latest access date: 03/22/2025). **C, D.** Readcount % of top 4 frequent Phyla: Proteobacteria (blue), Firmicutes (yellow), Ascomycota (green), Bacteroidetes (red), and others (gray). **E-F.** Firmicutes % in male and female animals. Data were analyzed by One Way ANOVA (ns = non- significant; *P < 0.05; ** P < 0.01).

## IV. DISCUSSION

Sch-PAH is the most common form of PAH worldwide^9^ and has recently been associated with gut and lung microbiome dysbiosis and significant lung microvascular apoptosis in severely remodeled vessels.^4,7^ Dysfunctional gut-lung microbiome composition can lead to increased levels of eATP, contributing to sustained tissue injury. Functional pathway analysis of lung metadata from the preclinical Sch-PH model indicated a reduction in aerobic respiration compared to controls, often indicative of cell stress via mitochondrial dysfunction, known to increase ATP secretion. *Ex vivo* quantification of ATP is often challenging due to its very short lifespan. Specifically, once ATP is released from injured cells or activated immune cells in response to pathogens, it can be either rapidly hydrolyzed by ectonucleotidases or activate purinergic receptors such as P2X7R.^40,41^ Whereas P2X7R is primarily recognized for its damage-associated role, it has also been implicated in regulating the gut microbiota homeostasis in mice, potentially through its known microbicidal activities.^23^ Previously, in experimental cercarial-driven experimental schistosomiasis using *Swiss* mice, reduced P2X7R expression and function were observed in mesenteric ECs and peritoneal macrophages,^16,42^ in contrast to elevated lung EC- P2X7R expression observed in Sch-PH. This data suggests that P2X7R expression may be differentially regulated depending on the organ, vascular bed, and/or disease stage. Specifically, endothelial P2X7R expression was upregulated only in the lung microvasculature, reaffirming its contribution to microvascular apoptosis in Sch-PH. Although P2X7R-mediated NLRP3 inflammasome has been implicated in PH, this pathway was not further evaluated in the Sch-PH model, leaving an important open gap for future research.^18^ Pharmacological inhibition of P2X7R, however, was evaluated using BBG. The inhibition of P2X7R significantly decreased vascular remodeling and RVSP in Sch-PH, although RVH was not ameliorated. This could potentially be due to the dose regimen or therapeutic time points of the BBG treatment or, more importantly, the direct effects of the *S. mansoni* egg antigens on the heart tissue itself; regardless, these results indicate the P2X7R may be a potential therapeutic target during the Sch-PH pathogenesis.

P2X7R co-localizes with the protein Cav-1 in healthy lung vasculature, where massive ATP efflux occurs in response to inflammation, infection, and tissue injury.^36–38^ Cav-1 is known as an anti-inflammatory and scaffolding protein crucial for pulmonary microvascular integrity, and its depletion via protein degradation and extracellular vesicle shedding is linked to vascular injury and the expansion of an apoptosis-resistant EC phenotype in experimental, preclinical, and human PAH.^7,11,43^ Moreover, Cav-1 expression has been previously implicated in the regulation of the anti-apoptotic IAP member survivin^39^, but its impact on other IAP family members was unknown. Therefore, reduced EC-Cav-1 may contribute to c-IAP2 depletion observed in egg- exposed mice, but the opposite seems not to be fully supported by the data from our new animal model. Inducible c-IAP2, but not constitutive c-IAP1, was observed to be critical for EC resistance to TNF-α-induced apoptosis.^44^ In PAH, TNF-α has been reported to reduce expression of Cav-1 but also of BMPRII, which is essential for lung EC homeostasis. Similarly, not only Cav-1 but also BMPRII expression is reduced in the lungs of Sch-PH animals,^7^ which may suggest a complex link between Cav-1/BMPRII and anti-apoptotic c-IAP2-mediated signaling in pathogen-induced PH. In line with decreased c-IAP2 expression observations, our data indicate increased circulating levels of c-IAP2 in the plasma of *S. mansoni* egg-exposed animals, suggesting that lung endothelial c-IAP2 may be shed along with or through similar mechanisms as lung EC-Cav-1 and could serve as a potential biomarker for disease severity.

Previous studies using a global c-IAP1/2 knockout model in a Rosa-creER^T2^ background (i.e., *R26- CreER^T^*^2^*; c-IAP1^-/-^; c-IAP2^fl/fl^*) revealed that systemic genetic deletion of c-IAP1 and c-IAP2 expression led to embryonic death, while conditional genetic ablation of c-IAP2 through *c-IAP2^fl/fl^* contributed to mild lung inflammation.^30^ Furthermore, c-IAP1 and c-IAP2 were also found to suppress caspase-8-dependent death *in vivo*, particularly in the intestines and liver,^30^ which may contribute to microbiome compositional or functional changes. Using *c-IAP1^-/-^; c-IAP2^fl/fl^* background under endothelial promoter expression control, we generated, validated, and induced the Sch-PH model, which revealed spontaneous increased RVH and RVSP and exacerbated remodeling at baseline and when exposed to IP/IV eggs. Specifically, genetic ablation of endothelial c-IAP2 expression resulted in increased lung microvascular remodeling in the homozygous c-IAP2 model compared to control animals. Moreover, rodent echocardiographic analysis highlighted early pulmonary vascular and RV changes associated with the development of PH or RV overload. Specifically, *S. mansoni* egg stimulation caused a significant reduction in PAT and the PAT/PET ratio but no changes in PET alone, suggesting increased pulmonary vascular resistance. RVFWT was significantly increased, confirming RVH, while fractional shortening, TAPSE, and other global cardiac performance parameters, including CO, EF, SV, and HR, showed no significant changes. These findings indicate that the endothelial expression of c-IAP2 plays a critical role in the development of Sch-PH. Monitoring trends in these parameters over time in egg-exposed animals alone or after pharmacological P2X7R inhibition could provide valuable insights into the progression or stabilization of pulmonary and RV function. Together, this study underscores the localized and early effects on the RV and pulmonary vasculature in the context of Sch-PH and highlights the relevance of endothelial c-IAP2 in driving these pathological changes.

Although recent work uncovered that gut-lung microbiome dysbiosis is a contributing factor to schistosomiasis-associated PH,^7^ the underlying mechanisms remained unclear. This study identifies a putative mechanistic link between lung microbiome dysbiosis, eATP accumulation, and P2X7R activation, culminating in c-IAP2 suppression and driving lung microvascular endothelial apoptosis. This represents, to the best of our knowledge, the first suggested connection between endothelial P2X7R and c-IAP2 in the context of PAH and microbiome dysbiosis. These findings offer new insight into how interorgan interactions may influence endothelial homeostasis in PH, a complex and understudied process. Moreover, the potential for sex-specific effects of lung microbiome dysbiosis on PAH onset and development remains poorly defined, underscoring the importance of further investigation. While metagenomic studies continue to expand our understanding of host-microbiome interactions in health and disease, current technical challenges - such as high data variability, limited reproducibility, and insufficient support for cross-model comparisons - highlight the need for supporting more extensive validation within and across systems to support the development of meaningful translational and biotherapeutic approaches.

Beyond the findings of this study and the comments above, it must be noted that this study had a few limitations. In terms of echocardiographs, for both IV injections of the *S. mansoni* eggs and raw data collection, the mice were under isoflurane-induced anesthesia. Isoflurane, a volatile anesthetic, has been observed to inhibit toll-like receptor 4 (TLR4)-NLRP3 inflammasome activation in microglia in diabetic mice.^45^ Previous studies have indicated that genetic ablation of TLR4 impaired the development of hypoxia-induced PH in mice.^30^ Since P2X7R is known to activate NLRP3, even though isoflurane’s effects in the lungs are unclear, this potential relationship should be noted. Furthermore, our endothelial-specific c-IAP2 flox model involves the systemic depletion of constitutive c-IAP1 expression, which did not appear to have significant effects on lung vascular homeostasis. Regardless of these limitations, our studies using the preclinical animal model of Sch-PH find for the first time that c-IAP2 expression is decreased in the lung tissue and increased in the circulation of *S. mansoni* egg-exposed mice. Data also showed that genetic ablation of endothelial c-IAP2 expression leads to spontaneous increases in microvessel thickness and area, suggesting that c-IAP2 expression and function may be critical for lung vascular homeostasis. Overall, these findings support the role of c-IAP2 as a novel therapeutic target and potential biomarker for Sch-PH **(Fig. 7)**. In conclusion, although the mechanisms supporting the survival of this abnormal lung EC phenotype remain in need of further studies, our data suggest that overactivation of microvascular P2X7R and impaired lung c-IAP2 expression may contribute to the expansion of this phenotype in Sch-PAH. Together, these findings offer new insights into the complex pathogenesis of Sch-PAH and highlight the importance of targeting c-IAP2 and P2X7R in promising future therapeutic strategies.

**Figure 7.**
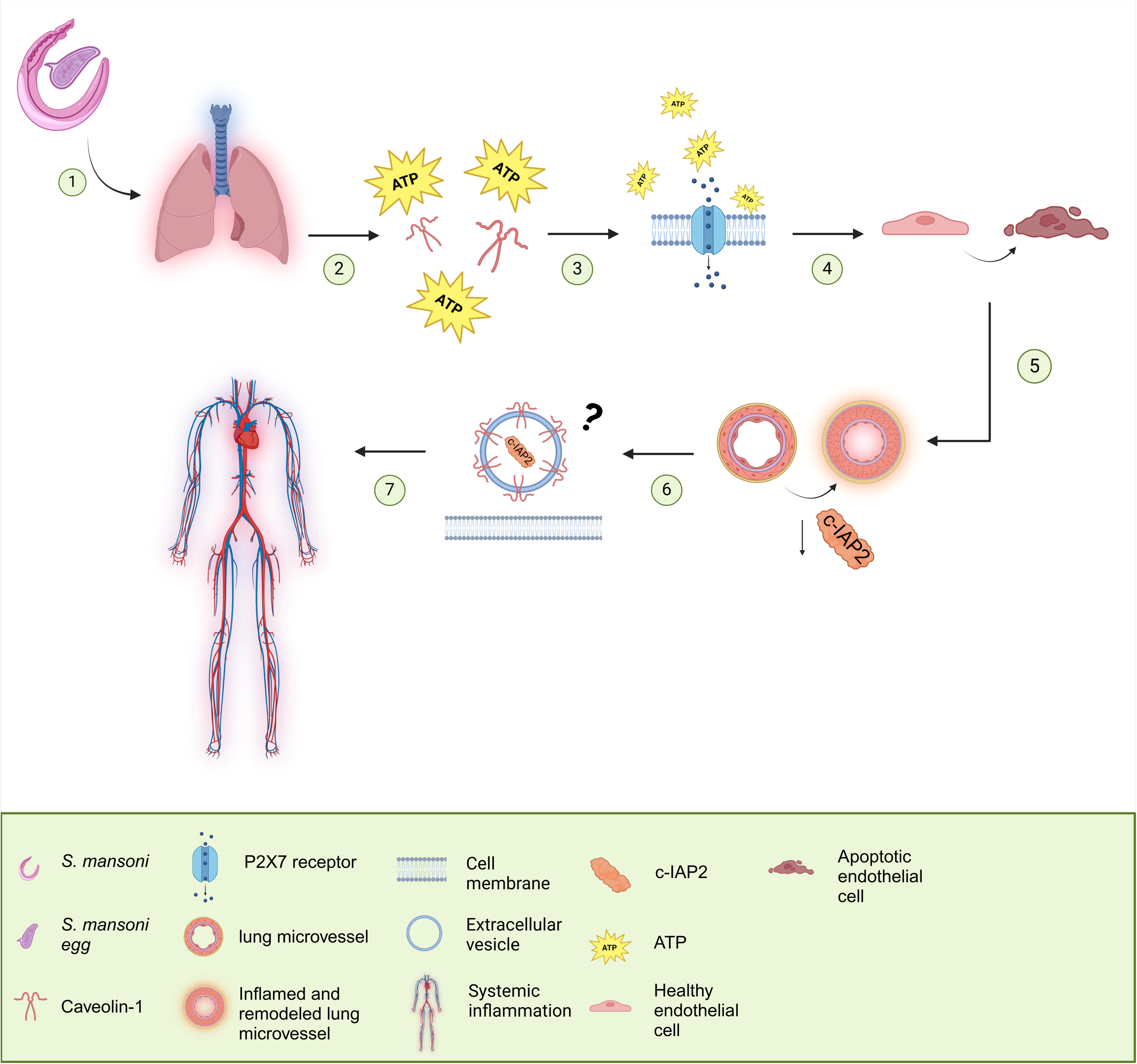
Representation of how lung endothelial c-IAP2 expression is lost in the circulatory system: Representation figure of *Schistosoma Mansoni* (*S. mansoni)* infection leading to lung microbiome dysbiosis (1). Lung dysbiosis and infection lead to increased adenosine triphosphate (ATP) shedding and Caveolin-1 (Cav-1) depletion (2). The increased ATP in the circulation and depleted Cav-1 lead to increased purinergic receptor family member P2X7 (P2X7R) activation (3). Increased P2X7R function leads to elevated apoptosis in endothelial cells (ECs) (4). Increased inflammation and apoptosis contribute to microvascular remodeling and loss of the inhibitors of apoptosis family member c-IAP2 (5). Lost c-IAP2 may be packaged into vesicles by Cav-1 and shed into the circulation, similar to Cav-1 (6); loss of c-IAP2 and Cav-1 combined with the overactivation of P2X7R leads to the propagation of Sch-PH (7). Created with BioRender.com

## ACKNOWLEDGEMENTS

We thank Genentech for kindly providing the frozen gametes of the *cIAP1^-/-^,cIAP2^fl/fl^* strain used to generate our novel endothelial-specific *Cdh5cre-ER^T^*^2^*;cIAP1^-/-^,cIAP2^fl/fl^*animal model. We also thank the Northwestern Transgenic and Targeted Mutagenesis Laboratory for recovering the *cIAP1^-/-^,cIAP2^fl/fl^* strain; the Lab technician MS Maricela Castellon for her technical work in animal husbandry and sample collection; UIC Research Resources Core (RRC), specifically the Mass Spectrometry Director, Dr. Hui Chen, for technical support in MS-LC, the Cardiovascular Core Director, Dr. Jiwang Chen and the Research Assistant Sam Lee for technical support in rodent hemodynamics/echocardiography, and the Histology Core Director Dr. Maria Sverdlov. Finally, infected *B*. *glabrata* snails were provided by the National Institute of Allergy and Infectious Diseases (NIAID) Schistosomiasis Resource Center for distribution through BEI Resources, NIAID, NIH.

## V. DISCLOSURES

*Cdh5cre-ER^T^*^2^*;cIAP1^-/-^,cIAP2^fl/fl^* animal model has been generated and validated under disclosure protection by the material transfer agreement (MTA) #106923 between UIC and Genentech. Therefore, part of the data is freely available, and part is available at reasonable request due to privacy agreement restrictions.

## VI. FUNDING SUPPORT

This work has been supported by the National Institutes of Health (NIH HL159037) and in part by the UIC Dept. of Anesthesiology and the UIC Health Equity Pilot Project (HEPP) CDA Award (SDO). Additionally, this project was supported in part by a 2022-2023 UIC Liberal Arts & Sciences Undergraduate Research Initiative (LASURI) scholarship, the 2024 UIC Honors Research Grant Award, and the 2024-2025 UIC Portal to Biomedical Research Careers (UIC PBRC) PREP Training grant (NIH/NIGMS 2R25 GM121212-06); and PVRI Travel Grant (EV). Additional support came from NIH/NIAID project AI177493 to DLW.

## Notes

### Competing Interest Statement

The authors have declared no competing interest.

