## Supplementary material for "Endothelial c-IAP2 Loss Amplifies P2X7 Receptor-Driven Inflammation and Worsens Infection-Associated Pulmonary Hypertension": Suppl Figs

**SUPPLEMENTARY FIGURES:**

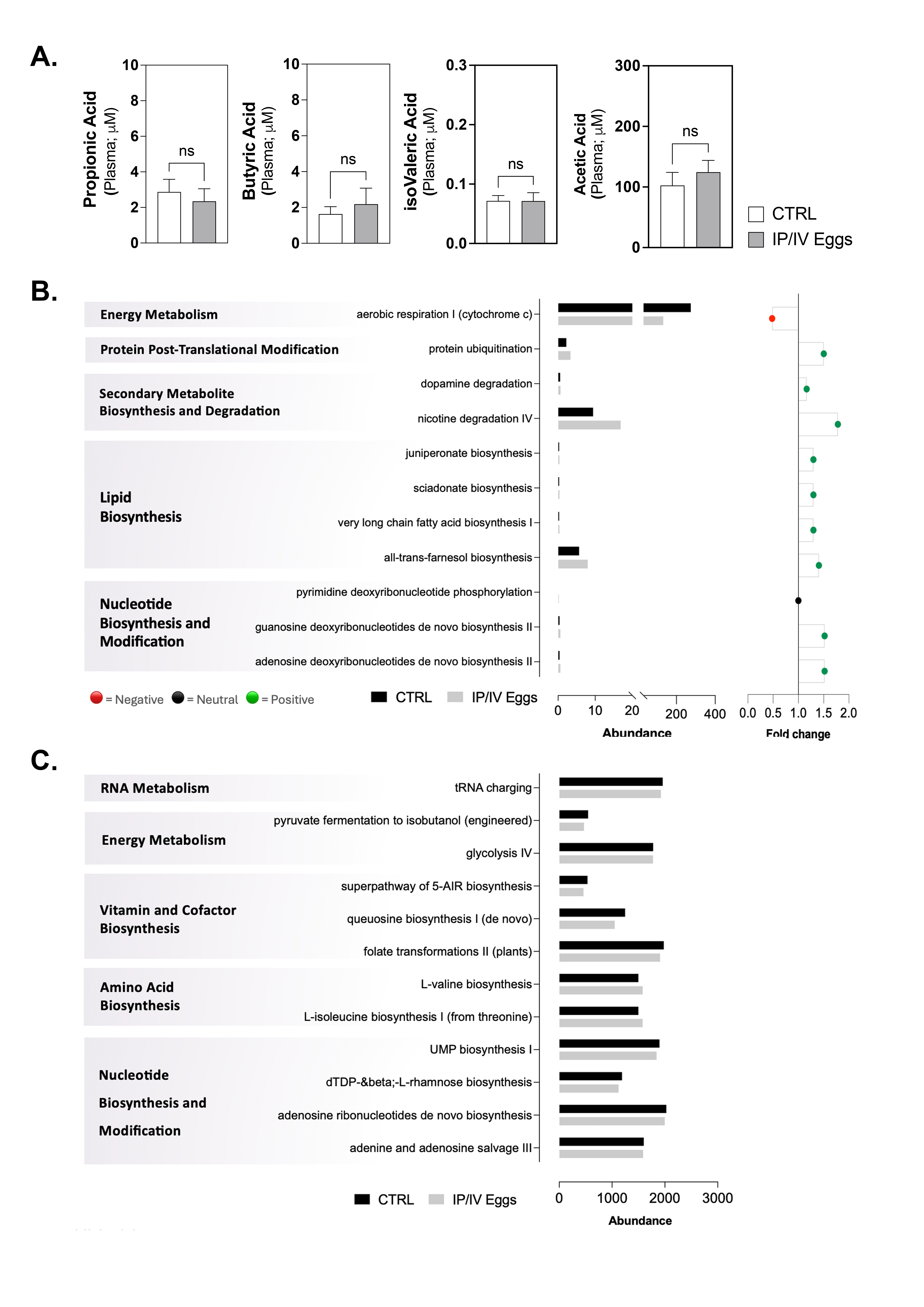

**Suppl. Figure 1: A.** Short-chain fatty acids (SCFAs) measured in plasma from vehicle control (CTRL) or mice exposed to intraperitoneal (IP) sensibilization using 240 *S. mansoni* eggs/gram of body weight (bw) followed by intravenous (IV) tail injection with 175 eggs/gram bw after two weeks (IP/IV Eggs). Metabolic pathway abundance was defined by the MetaCyc database using MinPath to determine the minimal pathway reconstruction based on gene families in the metadata from lung (**B**) or gut-derived **(C**) samples. After characterizing species composition, reads will be mapped to a custom annotated database (with bowtie2 + ChocoPhlAn) to identify gene families and taxonomy. Diamond and the UniRef90 database will be used to translate the search on any unmapped reads, identifying gene families. Next, gene family data and functional processes will be annotated into the MetaCyc database using MinPath. Normally distributed data was analyzed using a Student *t*-test (n = 8-10 animals/group; n = three different cultures; ns = non-significant).

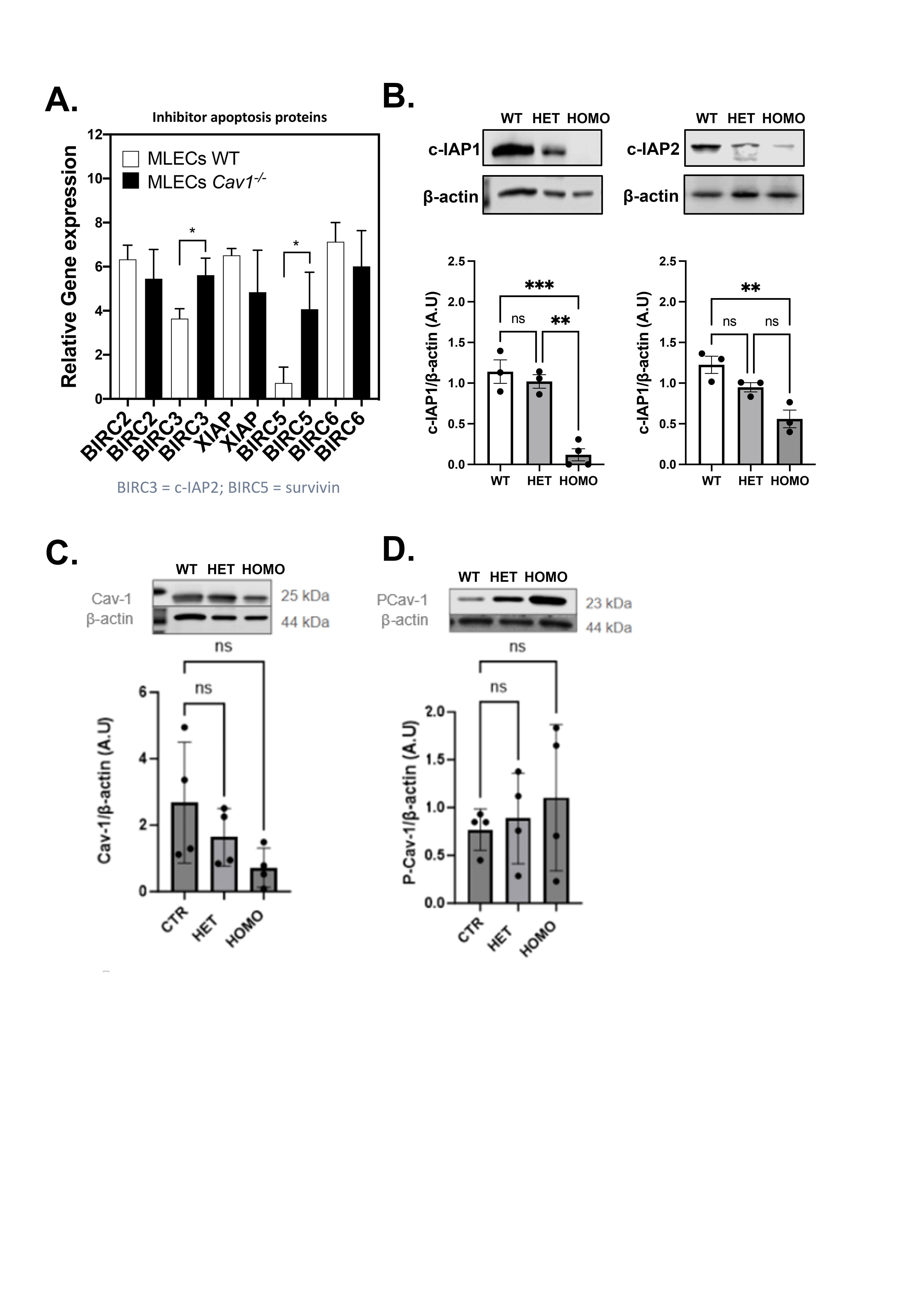

**Suppl. Figure 2: A.** Relative gene expression showing BIRC2 (c-IAP1), BIRC3 (c-IAP2), XIAP, BIRC5 (survivin), and BIRC6 (Bruce/Apollon) on wild-type (WT) versus *Cav1^-/-^* murine lung endothelial cells (MLECs). **B-D.** Western blot analysis of c-IAP1, c-IAP2, Cav-1, and Phosphorylated Cav-1 (PCav-1) in control (CTRL), heterozygous (HET), and homozygous (HOMO) mouse strain (*Cdh5creER^t2^; c-IAP1^+/-^:c-IAP2^+/fl^* and *Cdh5cre-ER^T2^;cIAP1^-/-^,cIAP2^fl/fl^*, respectively) (n = 3-6 animals/group; ns = non-significant; *P < 0.05; ** P < 0.01). = non-significant; *P < 0.05; ** P < 0.01; *** P < 0.001).
