## Supplementary material for "Endothelial c-IAP2 Loss Amplifies P2X7 Receptor-Driven Inflammation and Worsens Infection-Associated Pulmonary Hypertension": Suppl Material and Methods

**EXTENDED MATERIAL AND METHODS**

**2.4. Genotyping using Polymerase Chain Reaction:** Approximately 1 cm tail samples were collected from adult animals in a sterile environment and used to perform PCR analysis. PCR was performed via a REDExtract-N-AMP^TM^ Tissue PCR kit protocol (Sigma, Cat #XNAT-100Rxn)**.** For the tail digestion, a solution containing 50 µl of extraction solution and 12.5 µl of tissue preparation solution was mixed with individual tail samples and incubated at room temperature for 10 minutes. After the initial incubation, the samples were incubated at 95^º^C for 3 minutes before mixing them with 50 µl of neutralizing solution B. Different genotyping protocols were performed depending on the protocol specificities of the gene of interest. For identification of *Cdhr5*, forward primer (5’-GAT CGC TGC CAG GAT ATA CG-3’) and reverse primer (5’-AAT CGC CAT CTT CCA GCA G-3’) were used. After mixing each sample, the tubes were transferred to a thermal cycler (Applied Biosystems, 2720 Thermal Cycler) for the PCR reaction. Initial denaturation occurred at 94^º^C for 6 minutes, denaturation occurred at 94^º^C for 1 minute, annealing occurred at 55^º^C for 1 minute, extension occurred at 72^º^C for 1 minute and 30 seconds, and final extension occurred at 72^º^C for 5 minutes. 30-35 cycles of denaturation, annealing, and extension were completed before moving on to the final hold at 4^º^C. In parallel, *Birc3* PCR contained the same amount of reagent and tissue extracts as Cdhr5, except that three primers were used instead of two and had the same annealing and extension cycle. The primers used for *Birc3* were: 5’-GTG GTT TCC AAC GGC TTT G-3’; 5’-AAG TCT AGT CAC AGA GGC TCC AGT-3’, and 5’-GAT GGT GGC ACA TGC CTT TAA TCC-3’. The PCR products were then run in a 2-3% agarose gel electrophoresis, and subsequently, the gels were scanned with Li-Cor Odyssey CLx (Lincoln, NE).

**2.6. Experimental Rodent Echocardiography:** At day 0 (D0 - baseline) or day 21 (D21) after IV/IP PBS or Egg exposure, heterozygous and homozygous c-IAP2 mice were anesthetized using inhaled isoflurane (2.5% - 3%) and placed in the supine position on a heating pad. Then, individual animals were subjected to transthoracic echocardiography using Vevo F2 (VisualSonics Inc., Toronto, ON, Canada) and a UHF57x transducer. A rectal probe continuously monitored body temperature (T = 36.5-37.5 °C). The respiration rate was monitored and controlled by adjusting the depth of anesthesia. RV fractional area change (RVFAC) was measured via parasternal short-axis view at mid-papillary level. RV free wall thickness (RVFWT) was calculated during end-diastole in the parasternal short-axis mitral valve level, two-dimensional or parasternal long-axis RV outflow tract level, M-mode. In the short-axis view, a pulse-wave Doppler echo was used to record the pulmonary blood outflow at the aortic valve level to measure pulmonary acceleration time (PAT) and ejection time  (PET). Tricuspid annular plane systolic excursion (TAPSE) was measured in 2D M-mode echocardiograms from the apical 4-chamber view, positioning the cursor on the lateral tricuspid annulus near the free RV wall and aligning it as close as possible to the apex of the heart. Stroke volume (SV), fractional shortening (FS), ejection fraction (EF), and cardiac output (CO) were measured from the left ventricle (LV). The severity of pulmonary vascular remodeling (described below) and RVH (described above) was evaluated by histological analysis at the experimental endpoint.

**2.7. Histological analysis of Pulmonary Vascular Remodeling and IHC:** PFA-fixed, paraffin-embedded lung sections (5 μm) were used to evaluate protein expression (primary antibodies: CD31 (R&D systems; AF3628), c-IAP2 (R&D systems; MAB817), α-SMA (Invitrogen; 710487), and P2X7R (Protein Tech; APR-008) and for histological analysis. Samples were deparaffinized by serial exposure to xylene and ethanol. Then, antigen retrieval was performed for 5-10 min at ~ 120°C under a pressurized container using 1X sodium citrate as a buffer. For IHC, lung sections were washed with a 0.03% Triton 100x (Sigma-Aldrich, Cat # T8787) buffer diluted in 1X PBS for 2 x 5 minutes. Then, the slides were blocked using 10% goat serum or 5% BSA diluted in PBS (1 hour; room temperature), followed by overnight incubation with the primary antibody at 4^o^C (in the humidified chamber). After two periods of 5 minutes of washing, slides were incubated with secondary antibodies, rewashed (3 x 5 minutes), and mounted using mounting media containing DAPI. Then, fluorescent images were collected using an LSM880 confocal microscope (Carl Zeiss MicroImaging, Inc.). Fluorescent images were taken from randomized peripheral lung segments in micro- and macrovessels in each sample to quantify the expression of the proteins of interest. In addition, histological analysis of microvessel area and thickness was quantified in *Masson’s Trichrome-*stained sections. After staining, slides containing 1-2 sections were scanned using an Aperio brightfield automated microscope slide scanner (40X; Leica Aperio AT2). Digitalized images were used to determine the microvessel area and thickness (μm) in 10-20 microvessels/animal, i.e., vessels with a diameter smaller than 100 μm, using the ImageScope software 12.4.6 (Leica Biosystems). Briefly, microvessel wall thickness was obtained via the ImageScope ruler tool to measure the length across the thickest part of the vessel segment, defined as the space between the peripheral outer wall of the vasculature and the luminal boundary. Microvessel wall area was collected by tracing the contours of the peripheral outer wall and luminal boundary to obtain the total area and luminal area, respectively, and calculated by subtracting the luminal area from the total area.^3^

**2.9. Competitive Enzyme-Linked Immunosorbent Assay:** Competitive Enzyme-Linked Immunosorbent Assay (ELISA) was carried out following the protocol from the mouse baculoviral IAP repeat-containing protein 3 (BIRC3; c-IAP2) ELISA Kit (MyBioSource, Cat #MBS7252940). The ideal dilution of plasma samples from control or egg-exposed mice (i.e., 1:4) was standardized using 1x DPBS before ELISA. All ELISA kit components were brought to room temperature before 100 µl of standards (A-F), and designated samples were added to the appropriate wells in duplicate amounts; blanks containing only 1x DPBS were also used. 50 µl of conjugate was added to each well and mixed thoroughly before the plate was covered and incubated for 1 hour at 37 °C. After 1 hour, the plate was manually washed by aspirating the wells and filling each well entirely with 1x wash solution. The manual washing was completed 5x before the plate was inverted and blotted against absorbent paper. 50 µl substrate A and 50 µl substrate B were added to all wells and incubated for 15 minutes at 37 °C before 50 µl of stop solution was added to all wells. The optical density was then measured at 450 nm in a microplate reader (Accuris Smartreader 96), and the results were plotted on GraphPad Prism.

**3.2. Sample Preparation and Western Blot:** Frozen lung tissue and cultured ECs were fully homogenized using cold radioimmunoprecipitation assay (RIPA) buffer containing 1% protease and 0.1% phosphatase inhibitor cocktail. After 20 min of incubation at 4 ^o^C, homogenates were centrifuged at 12,000 x g (20 min at 4^o^C), and the supernatant was collected for protein measurement. Standard bicinchoninic acid (BCA) protein assay was used to determine the protein concentration of each lung sample by colorimetry using a microplate reader (Benchmark Scientific MR9600-T SmartReader™ 96 Plate). Then, 10-30 μg of lung tissue lysates were diluted in Laemmli Sample Buffer (4X) with ß-mercaptoethanol and boiled for 10 min at 95^o^C. Before loading the samples on a gradient SDS-PAGE gel (8%–12%), tubes were centrifuged for 2.5 min at 16,200 x g. After running the samples, proteins were transferred to nitrocellulose membranes, and the transfer was assessed by the absence of ladder markers in the gel, simultaneously detecting a continuous gradient of Ponceau Rouge staining over the membranes. Then, membranes were washed with TBS-Tween 1X for 5 min twice and blocked using 5% milk or BSA (according to antibody datasheet instructions) for 1 hour at room temperature, followed by primary antibody incubation (overnight at 4 ^o^C or 2-3 hours at 37 ^o^C). After washing (2 x 5 min; 1 x 15 min), the membranes were incubated for 1 hour with the specific secondary HRP-conjugated antibody, washed again, and then detected using an ECL kit (Amersham, Piscataway, NJ). Membranes were scanned with Li-Cor Odyssey CLx (Lincoln, NE), and data were analyzed and normalized to β-actin or GAPDH loading controls using ImageJ software (<https://imagej.nih.gov/ij/>).

**3.5. Reagents and Antibodies:** REDExtract-N-AMP^TM^ Tissue PCR kit protocol (Sigma, Cat #XNAT-100Rxn), baculoviral IAP repeat-containing protein 3 (BIRC3; c-IAP2) ELISA Kit (MyBioSource, Cat #MBS7252940), DPBS, ECL kit (Super Signal West Pico PLUS; REF: 34580; Thermo Scientific), Bio-Rad Protein Assay Kit II (Cat No. 5000002; Bio-Rad), Radioimmunoprecipitation assay (RIPA; Cat No. J63306.AK; Thermo Scientific), APC Annexin V Apoptosis Detection kit with PI (Cat No. 640932; Biolegend), Trypsin-EDTA 1× (Cat No. 15400-054; Gibco), Staurosporine (STS; Cat No. 1285; Tocris), A740003 (Cat. No. 3701, Tocris), TNF-ɑ (Cat No. GF023; Millipore), INF-Ɣ(Cat No. IF002; Millipore), ATP (Cat No. A2383; Sigma Aldrich), Brilliant Blue G (BBG; P2X7R pharmacological inhibitor; Cat No. BB0770; Sigma), Endothelial basal medium-2 (EBM-2; Cat No. CC-3156; Lonza), MV SingleQuots (Cat No. CC-4147; Lonza), SingleQuots (Cat No. CC-4176; Lonza), Corn oil (Cat No. C8267; Sigma Aldrich), Tamoxifen (Cat No. J63509.03; Thermo Scientific). Rabbit polyclonal anti-α-SMA was acquired from Abcam (Boston, MA, USA). Rabbit polyclonal anti-GAPDH was acquired from Santa Cruz Biotechnology (Santa Cruz, CA, USA). Alexa-Fluor 488 and 555-conjugated goat anti-mouse and anti-rabbit IgG were purchased from Life Technologies (Grand Island, NY, USA). Anti-mouse and anti-rabbit HRP-conjugated IgG were purchased from Cell Signaling Technology (Danvers, MA, USA) or Kierkegaard & Perry Laboratories (Gaithersburg, MD). RIPA buffer, protease and phosphatase inhibitor cocktail, collagenase type I, PFA, sodium citrate, heparin, and sucrose were purchased from SIGMA Chemical Co. (St. Louis, MO, USA). Mounting media with DAPI (VectaShield) was obtained from Vector (Burlingame, CA, USA). The batches 03614, 03577, and 03228 from Sm-p40 were acquired from Biomatik (Wilmington, Delaware, USA). Stock solutions were prepared in 100% dimethyl sulphoxide (DMSO) or sterile PBS and diluted daily in sterile PBS or cell medium for *in vitro* treatments. The highest final concentration of the solvent was 0.1% (v/v) and did not affect the experiments. The PCR primers were purchased from Integrated DNA Technologies, Inc. (Coralville, Iowa, USA).
